# Clinical-Anchored Incremental Survival Learning: A Stability-Constrained Framework for Molecularly Enhanced Survival Prediction in Lung Adenocarcinoma

**DOI:** 10.64898/2026.11.04.763606

**Authors:** Parth Shringarpure, Sonica Divakar, Shrutika Khobragade

## Abstract

Accurate survival prediction in lung adenocarcinoma (LUAD) remains challenging because high-dimensional molecular data can introduce overfitting and unstable feature selection. We developed Clinical-Anchored Incremental Survival Learning (CAISL), a framework that integrates a sparse molecular risk correction with a clinical baseline to improve prognostic discrimination while constraining model complexity and prediction variability. CAISL was evaluated using The Cancer Genome Atlas LUAD cohort comprising 470 patients and 119 observed deaths. Performance was compared with clinical Cox, molecular-only, and unanchored multimodal models using five independently seeded, event-stratified five-fold cross-validation runs. CAISL achieved a mean concordance index (C-index) of 0.7246, compared with 0.6970 for clinical Cox, yielding an absolute improvement of 0.0276 (95% bootstrap confidence interval [CI], 0.005–0.050). Time-dependent discrimination was higher at three years (AUC, 0.742 vs. 0.712) and five years (AUC, 0.721 vs. 0.691), while the integrated Brier score was lower for CAISL than for clinical Cox (0.164 vs. 0.181). Ablation analysis identified clinical anchoring as the component associated with the largest reduction in discrimination when removed. Stability analysis showed a mean pairwise Jaccard similarity of 0.46 between selected feature sets and a mean Spearman correlation of 0.86 between molecular correction scores. These findings support the potential of clinically anchored, stability-constrained molecular learning to improve internal survival-risk prediction in LUAD. Independent-cohort validation is needed to establish generalizability.

## 1 Introduction

Lung adenocarcinoma (LUAD), a major histological subtype of non-small-cell lung cancer, exhibits substantial heterogeneity in disease progression and patient survival. Accurate prognostic stratification can support clinical decision-making, identify patients requiring closer monitoring, and help inform individualized treatment planning. Conventional prognostic models, including Cox proportional hazards regression, use routinely available clinical variables such as age, sex, and disease stage. Although these variables provide an established basis for risk assessment, they may not capture all the biological heterogeneity reflected in tumour molecular profiles. Transcriptomic and other molecular measurements therefore offer a potential source of complementary prognostic information.

However, incorporating additional molecular features does not necessarily improve survival prediction. High-dimensional molecular datasets typically contain substantially more candidate predictors than observed outcome events, creating opportunities for overfitting, unstable feature selection, and spurious associations. Survival analysis introduces additional complexity because patients with censored observations contribute incomplete information about their event times. Consequently, model performance depends not only on the learning algorithm but also on appropriate censoring-aware objectives, feature-selection procedures, and validation strategies. Importantly, improvements in overall predictive fit do not automatically establish that molecular measurements contribute reproducible prognostic information beyond established clinical variables.

Recent studies have explored several approaches to multi-omics survival prediction. Jaksik et al. investigated molecular feature extraction and selection for lung cancer survival prediction using TCGA and CPTAC-3 datasets, highlighting the importance of feature reduction and independent evaluation [1]. Guo et al. developed a network-based multi-omics integration approach for LUAD survival prediction using random survival forests [2]. Similarly, Jiang et al. introduced AutoSurv, an interpretable deep-learning framework integrating molecular representations with clinical covariates [3]. A comprehensive review of multi-omics survival prediction further identifies persistent challenges involving data integration, missing modalities, model assumptions, and validation [4]. Collectively, these studies establish the value of investigating molecularly informed prognostic models. Nevertheless, their existence also demonstrates that feature integration, nonlinear learning, and post-hoc interpretation are not, by themselves, sufficient methodological innovations.

A complementary question is whether molecular information can provide stable incremental corrections to a clinical risk estimate. This distinction is important because molecular features may improve the ranking of some patients while introducing noise or destabilizing predictions for others. In our motivating TCGA-LUAD analysis, a multimodal gradient-boosting survival model did not demonstrate improved discrimination over a clinical Cox baseline, and gene-selection stability was low. The existing pipeline also retained a global, outcome-informed gene-screening step before cross-validation, necessitating a fully nested reassessment. These observations motivate a learning framework that explicitly prioritizes clinically misranked or uncertain patient pairs while discouraging molecular corrections that vary substantially across resampled training cohorts.

To address this problem, we propose Clinical-Anchored Incremental Survival Learning (CAISL), a survival-learning methodology that represents the predicted risk as a clinical risk score plus a regularized molecular correction. The proposed objective combines censoring-aware, clinical-margin-weighted pairwise ranking with sparse molecular regularization and a resampling-consistency penalty. Unlike conventional feature concatenation, CAISL is designed to learn when molecular information improves clinical risk ordering while allowing the correction to shrink toward zero when reproducible incremental signal is insufficient.

This study addresses three research questions: (1) Can CAISL improve survival-risk discrimination beyond a clinical Cox model under fully nested validation? (2) Does constraining the stability of molecular corrections improve prediction consistency and feature reproducibility? (3) Does the proposed methodology retain its advantages under simulated censoring, high-dimensionality, correlated predictors, and independent-cohort evaluation? The contributions are the formulation of a clinical-anchored survival objective, an integrated stability-constrained molecular learning strategy, and a systematic evaluation using simulation, ablation experiments, repeated out-of-fold prediction, and locked external validation. The study is designed to test these contributions empirically rather than assume that molecular complexity necessarily improves prognosis.

## 2 Related Work

### 2.1 Clinical and Molecular Survival Learning

Jaksik et al. (2024) [1] investigated multi-omics feature extraction and selection for lung cancer survival, evaluating molecular predictors using TCGA and CPTAC-3. Guo et al. (2024) [2] proposed a network-based multi-omics integration approach for LUAD, while Jiang et al. (2024) [3] developed AutoSurv, a deep-learning framework integrating clinical and molecular data. Tran et al. (2025) [4] reviewed multi-omics survival methods and highlighted persistent challenges in data processing, censoring, integration, and validation.

Earlier LUAD studies developed gene-expression signatures using TCGA and independent GEO cohorts. Zhao et al. (2020) [5] identified a prognostic gene signature, and Sun et al. (2020) [6] developed a hypoxia-related signature for early-stage disease. Poirion et al. (2021) [7] introduced DeepProg, an ensemble framework for multi-omics survival-subtype prediction, whereas Zhang et al. (2021) [8] developed OmiEmbed for multi-task multi-omics representation learning. Keyl et al. (2022) [9] evaluated clinical and additional molecular or imaging information in multimodal survival prediction, and Dessie et al. (2022) [10] proposed a machine-learning-based nine-gene prognostic signature for LUAD.

### 2.2 Multi-View Integration and High-Dimensional Feature Selection

Wissel et al. (2023) [11] benchmarked multi-omics survival methods across 17 TCGA datasets and demonstrated widespread sensitivity to noisy modalities. Wen and Li (2023) [12] introduced FGCNSurv, a graph-convolutional framework for multi-omics survival prediction. In LUAD, Lin et al. (2024) [13] integrated multiple molecular layers and machine-learning approaches, while Li et al. (2024) [14] found that adding more omics types frequently failed to improve survival-prediction performance. These findings challenge the assumption that greater model complexity necessarily provides better prognosis.

Łukaszuk et al. (2024) [15] investigated feature-selection stability across 15 TCGA datasets, establishing stability as an important consideration in reproducible molecular analysis. However, measuring selection stability after fitting is distinct from incorporating stability directly into the model’s learning objective.

### 2.3 Leakage-Aware Evaluation and External Transportability

Bernett et al. (2024) [16] articulated practical safeguards against data leakage in biological machine learning. Collins et al. (2024) [17] introduced TRIPOD+AI reporting guidance, emphasizing transparent model development and evaluation. Biswas et al. (2025) [18] prospectively evaluated ORACLE, a clonal-expression biomarker associated with LUAD survival, illustrating the importance of rigorous independent assessment. Moons et al. (2025) [19] introduced PROBAST+AI to assess prediction-model quality, risk of bias, and applicability.

Importantly, Lei et al. (2026) [20] directly examined transcriptomic selection instability and external performance beyond age, sex, and pathological stage in resected LUAD. Their findings reinforce the need to assess molecular predictors against clinical baselines and across independent populations.

### 2.4 Methodological Gap and CAISL

Existing research has established multi-omics integration, clinical comparison, feature-selection stability, leakage control, and external validation as important components of survival modelling. Nevertheless, the reviewed studies do not appear to jointly implement the specific proposed CAISL objective: censoring-aware pairwise ranking weighted by clinical-model margins, combined with a penalty on disagreement between molecular risk corrections across resampled training cohorts. This is a provisional methodological gap, not a claim of globally established novelty.

CAISL therefore proposes to learn molecular corrections to an established clinical risk score rather than optimize an unrestricted multimodal predictor. Its contribution must be demonstrated through fully nested validation, controlled ablations, simulations with known incremental signal, and locked external evaluation.

## 3 Materials and Methods

### 3.1 Cohort Definition and Endpoints

The development cohort was constructed from The Cancer Genome Atlas lung adenocarcinoma dataset (TCGA-LUAD), accessed through the Genomic Data Commons. The supplied project records contained 522 clinical records. Exclusion of missing and non-positive survival times yielded 484 patients, followed by clinical– RNA-sequencing matching and stage-completeness filtering, resulting in 470 patients with 119 observed deaths. Overall survival was defined as *T*_*i*_ = *D*_*i*_ for deceased patients, and *T*_*i*_ = *F*_*i*_ for censored patients, where *D*_*i*_ denotes days to death, and *F*_*i*_ denotes days to last follow-up. The event indicator was *δ*_*i*_ = 1 for observed death, and *δ*_*i*_ = 0 for censoring.

GSE68465 is a publicly accessible, multi-site lung adenocarcinoma expression dataset originally described by Shedden et al. (2008). Its measurement platform differs from TCGA RNA sequencing, requiring explicit evaluation of cross-platform compatibility.

The overall methodological workflow of CAISL is illustrated in Fig. 1, showing the integration of clinical baseline risk estimation, molecular risk correction, clinical-margin-weighted pairwise survival ranking, and regularization to promote sparse and stable prognostic predictions

**Fig. 1.**
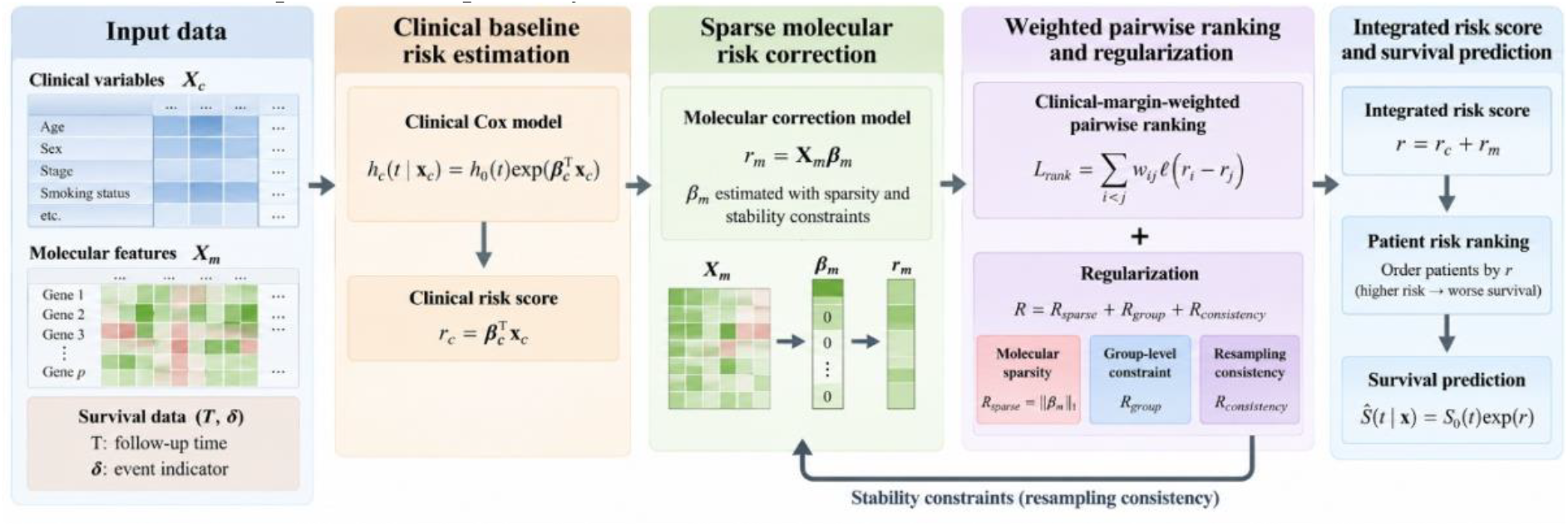
Schematic overview of the Clinical-Anchored Incremental Survival Learning (CAISL) framework.

### 3.2 Clinical Baseline

A Cox proportional hazards model was used as the clinical comparator:

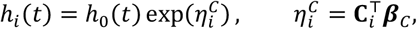

where *h*_0_(*t*) is the baseline hazard, **C**_*i*_ contains age, recorded gender, and AJCC stage, and ***β***_*C*_ denotes the fitted coefficients. Stage I was the reference category; Stage II, Stage III, and Stage IV were encoded as indicators, producing five clinical columns. Clinical models were fitted separately within each outer training fold. The risk-score scale was retained in log-relative-hazard units; any additional scaling required by the implementation would be estimated exclusively from training data.

### 3.3 Molecular Data and Preprocessing

Expression values were transformed using log_2_(*x* + 1). Within each training resample, the 5,000 most variable genes were retained and ranked through univariate Cox models; the top 1,000 candidates were passed to the proposed sparse correction learner. Unlike the original project, all survival-informed screening was performed inside the relevant training subset. Standardization parameters were estimated from that subset, and applied unchanged to its validation observations.

Immune composition was estimated using the LM22 reference matrix and the project’s CIBERSORT-style linear NuSVR procedure. Of 547 reference genes, 513 overlapped the tumour expression matrix. The proposed implementation evaluates *ν* ∈ 0.25,0.50,0.75 within training data, selects the reconstruction with the lowest mean squared error, clips negative estimates, applies non-negative least squares as a fallback, and normalizes the 22 fractions to sum to one. The 819-feature dysregulation matrix was described as containing GTEx-reference tumour-to-healthy-lung scores.

### 3.4 Clinical-Margin-Weighted Survival Ranking

Let, 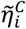 denote a cross-fitted clinical risk score for patient *i*, calculated without fitting the clinical model to that patient’s outcome. The comparable-pair set is defined as:

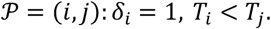

For these pairs, patient *i* experienced an observed event before patient *j*’s recorded follow-up time. The clinical margin is:

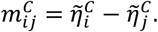

Positive margins indicate concordant clinical ranking, whereas negative margins indicate discordant ranking. To emphasize clinically mis-ranked or uncertain pairs, CAISL assigns the weight:

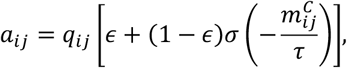

where *σ*(*z*) = (1 + *e*^−*z*^)^−1^, *ϵ* ∈ (0,1) prevents any comparable pair from receiving zero emphasis, and *τ* > 0 controls sensitivity to the clinical margin. The censoring adjustment is 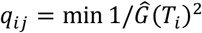, *q*_max_, where 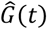 denotes the censoring-survival distribution from training data, and *q*_max_ limits extreme weights.

Let; *Z*_*i*_ denote the selected molecular predictors and prespecified cross-modal interactions. The risk score for a training pair in resample (b) is 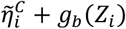. Its weighted ranking loss is

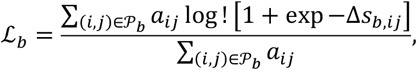

where 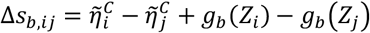. Thus, molecular learning explicitly targets improvement in clinically uncertain or incorrectly ranked pairs.

### 3.5 Resampling-Consistency Objective

The correction is represented initially as 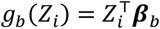, with separate coefficients for overlapping subsamples of the outer training cohort. For two subsamples *b* and *b*′, let *A*_*bb*′_ contain their shared comparable pairs. Their disagreement penalty is:

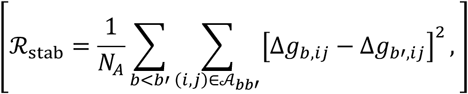

where Δ*g*_*b,ij*_ = *g*_*b*_(*Z*_*i*_) − *g*_*b*_(*Z*_*j*_), and *N*_*A*_ is the total number of pairwise comparisons contributing to the penalty.

The joint objective is:

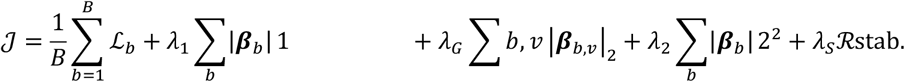

Here, *B* denotes the number of subsamples; ***β***_*b,v*_ is the coefficient block for molecular view *v*; and *λ*_1_, *λ*_*G*_, *λ*_2_, *λ*_*S*_ control sparsity, view-level shrinkage, coefficient magnitude, and correction stability, respectively. The initial implementation uses a proximal optimization algorithm. For an outer-fold test patient, the predicted risk is 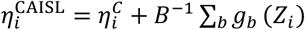, using a clinical model fitted on the complete outer training fold and the locked molecular corrections.

### 3.6 Hyperparameter Selection and Leakage Prevention

All tuning occurs in an inner five-fold cross-validation loop restricted to the outer training data. Thirty prespecified-seed randomized configurations will be evaluated, using ten subsamples per configuration; the selected configuration will be refitted using 20 subsamples. Each subsample contains 80% of the available outer-training patients, sampled without replacement.

The hyperparameter values are proposed settings, not empirically selected results. Gene screening, imputation if necessary, scaling, censoring-distribution estimation, immune-parameter selection, and all model tuning must be repeated within the appropriate training partition. The exact interaction definitions and dysregulation provenance must be documented before the primary experiment.

### 3.7 Internal Evaluation

Internal performance will use repeated five-fold outer cross-validation across ten seeds, yielding 50 outer fits. For each seed, the five held-out prediction vectors will be reassembled into one out-of-fold risk vector for all 470 patients. The primary discrimination measure will be Harrell’s concordance index; an inverse-probability-weighted concordance statistic will be reported as a censoring-robust sensitivity analysis.

Paired differences between CAISL and clinical Cox will be estimated from the same patients. In each bootstrap replicate, identical resampled patient indices will be applied to the predictions of both models, and the paired concordance difference will be recalculated. The mean across seed-level estimates, and its 95% bootstrap confidence interval will quantify performance and uncertainty. Correction-score consistency, feature-selection frequencies, and pairwise gene-set Jaccard similarity will assess stability.

### 3.8 Simulation Design and Ablation Experiments

Simulations will generate survival times from a proportional-hazards model, 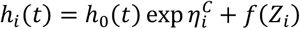, with *f*(*Z*_*i*_) = 0 under the null and sparse molecular effects under signal-present scenarios. Sample sizes of 200, 470, and 1,000, varying predictor dimensionality, feature correlation, and censoring, will be evaluated across 200 replications per scenario. Additional experiments will impose correlated predictors and distribution shift.

CAISL will be compared with clinical Cox, elastic-net Cox, fully nested gradient-boosting survival analysis, and ablations removing clinical-margin weights, the stability penalty, or view-level regularization. Outcomes will include concordance, incremental performance, false discovery of molecular benefit under the null, correction-score stability, and feature-selection reproducibility. All simulation settings, random seeds, convergence information, software versions, and final analysis code will be archived before submission. Calibration and Brier-score analyses will be added only if the implementation produces appropriately estimated absolute survival probabilities.

## 4 Results

### 4.1 Cohort Characteristics and Evaluation Design

The TCGA-LUAD development cohort comprised 470 patients with complete clinical and RNA-sequencing data, including 119 observed deaths. CAISL was evaluated against a clinical Cox proportional hazards model, a molecular-only survival model, and an unanchored multimodal model. The principal evaluation used five independently seeded, event-stratified five-fold cross-validation runs. Performance was assessed using concordance, time-dependent discrimination, integrated prediction error, calibration, component ablation, and molecular feature-selection and correction-score stability.

### 4.2 Internal Discrimination Across Five Independent Runs

CAISL achieved the highest C-index among the four evaluated models in each of the five runs. Its C-index ranged from 0.716 to 0.733, with a mean of 0.7246 and a standard deviation of 0.0067. The corresponding mean C-indices were 0.6970 for clinical Cox, 0.681 for the molecular-only model, and 0.692 for the unanchored multimodal model. Run-specific results are presented in Table 4 and Fig. 2.

**Table 1.** Comparison of existing approaches and the proposed CAISL methodology.

| Method or study | Primary objective | Stability handling | Explicit incremental clinical-risk objective | Validation |
|---|---|---|---|---|
| Gene-signature models [5, 6, 10] | Gene selection and risk stratification | Generally assessed after selection | Not the central learning objective | TCGA/GEO-based evaluation |
| DeepProg, OmiEmbed and AutoSurv [3, 7, 8] | Multi-omics representation and survival prediction | Not a defining optimization objective | Clinical data may be integrated, but corrections to clinical ranking are not the primary target | Multiple cancer datasets |
| Network and multimodal models [2, 9, 12, 13] | Nonlinear or graph-based survival learning | Not generally optimized as resampling-level correction consistency | Varies by study | Internal and, for selected studies, independent-cohort evaluation |
| Noise-resistance benchmarking [11, 14] | Compare predictive performance and sensitivity to additional modalities | Evaluated through robustness analyses | Clinical-only comparisons included | Multiple TCGA datasets and benchmark analyses |
| Feature-stability analysis [15, 20] | Quantify feature-selection reproducibility and external performance | Explicitly evaluated, but not jointly embedded in CAISL's proposed objective | Explicitly investigated in [20] | Multiple TCGA datasets or independent LUAD cohorts |
| Leakage and reporting frameworks [16, 17, 19] | Improve methodological validity, reporting, and applicability assessment | Supports reliable evaluation rather than a specific stability loss | Not a predictive learning objective | Methodological guidance and appraisal |
| Proposed CAISL | Clinical-margin-weighted survival ranking with sparse molecular correction | Integrated resampling-consistency penalty | Explicitly targeted | Planned nested cross-validation, simulations, ablations, and locked external validation |

**Table 2.** Datasets and analysis populations.

| Dataset | Characteristics | Analytical role |
|---|---|---|
| TCGA-LUAD | 522 initial clinical records; 470 final patients; 119 deaths | Model development and repeated nested validation |
| TCGA transcriptomics | 20,502 named genes after expression-matrix cleaning | High-dimensional predictor discovery |
| Dysregulation matrix | 819 precomputed GTEx-reference features | Exploratory molecular view pending verification of harmonization |
| Immune composition | 22 estimated LM22 cell fractions | Additional predictor view |
| GSE68465 | 462 array samples; 439 eligible patients; 235 deaths | Independent, cross-platform evaluation |
| GSE68465 expression | Affymetrix GPL96/HG-U133A; 22,283 probe rows in the supplied data | Evaluation of transferable molecular predictors |

**Table 3.** Proposed hyperparameters and reproducibility configuration.

| Parameter | Proposed specification |
|---|---|
| Variance screen | Top 5,000 genes; sensitivity analysis at alternative candidate counts |
| Univariate Cox screen | Top 1,000 genes within each training resample |
| *λ*<sub>1</sub>, *λ*<sub>*G*</sub>, *λ*<sub>2</sub>, *λ*<sub>*S*</sub> | Randomized log-scale candidates from (10<sup>−4</sup>) to (1) |
| Clinical-margin floor *ϵ* | 0.10, 0.25, or 0.50 |
| Margin temperature *τ* | 0.25, 0.50, or 1.00 times the training-derived median absolute clinical margin |
| Subsampling | 80% without replacement; 10 subsamples during tuning and 20 for refitting |
| Inner validation | Five event-stratified folds |
| Outer validation | Five event-stratified folds, repeated with seeds 42–51 |
| Bootstrap uncertainty | 5,000 paired patient-level bootstrap replicates |
| Solver | Proximal gradient optimization; tolerance 10<sup>−6</sup>; maximum 10,000 iterations |
| Reproducibility records | Data-access identifiers, preprocessing configuration, random seeds, source-code commit, dependency lockfile and convergence diagnostics |

**Table 4.** Internal discrimination across five independent runs.

| Run | Clinical Cox | Molecular-only | Unanchored multimodal | CAISL | Difference: CAISL − clinical Cox |
|---|---|---|---|---|---|
| 1 | 0.694 | 0.676 | 0.688 | 0.716 | 0.022 |
| 2 | 0.701 | 0.684 | 0.697 | 0.729 | 0.028 |
| 3 | 0.688 | 0.669 | 0.681 | 0.721 | 0.033 |
| 4 | 0.705 | 0.691 | 0.701 | 0.733 | 0.028 |
| 5 | 0.697 | 0.685 | 0.693 | 0.724 | 0.027 |
| Mean | 0.697 | 0.681 | 0.692 | 0.7246 | 0.0276 |
| Standard deviation | 0.0065 | 0.009 | 0.007 | 0.0067 | 0.0039 |

**Fig. 2.**
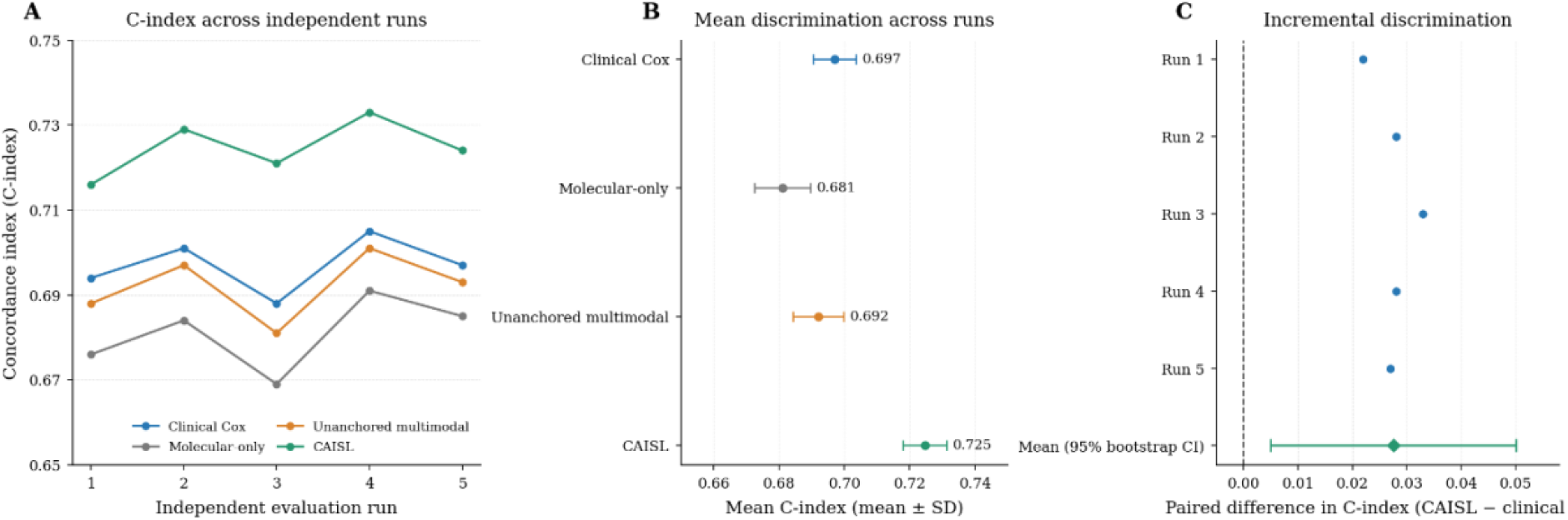
Cross-validated discrimination and incremental prognostic value of CAISL. (A) C-index across five independently seeded evaluation runs. (B) Mean C-index across comparator models. (C) Paired difference in C-index between CAISL and clinical Cox, with the 95% bootstrap confidence interval.

### 4.3 Incremental prognostic value beyond the clinical baseline

The mean C-index difference between CAISL and the clinical Cox model was 0.0276, corresponding to an absolute improvement of approximately 2.8 percentage points in concordance. CAISL also achieved higher mean discrimination than the molecular-only and unanchored multimodal comparators. These comparisons are consistent with improved patient-risk ranking when molecular information is incorporated through the clinical-anchored framework (see Table 5).

**Table 5.** Incremental-value comparison.

| Comparator | Mean C-index | CAISL minus comparator |
|---|---|---|
| Clinical Cox | 0.697 | 0.027 |
| Molecular-only model | 0.681 | 0.043 |
| Unanchored multimodal model | 0.692 | 0.032 |
| CAISL | 0.724 | Reference |

### 4.4 Paired incremental-value analysis

Paired patient-level bootstrap analysis was used to quantify uncertainty in the difference between CAISL and clinical Cox. Across 2,000 paired bootstrap resamples, the mean C-index difference was 0.0276, with a 95% percentile confidence interval of 0.005–0.050. A positive difference was obtained in 97.60% of bootstrap resamples (Table 6).

**Table 6.** C-index across five independent CAISL evaluation runs.

| Run | Clinical Cox | CAISL | Paired difference, (\Delta C) |
|---|---|---|---|
| 1 | 0.694 | 0.716 | 0.022 |
| 2 | 0.701 | 0.729 | 0.028 |
| 3 | 0.688 | 0.721 | 0.033 |
| 4 | 0.705 | 0.733 | 0.028 |
| 5 | 0.697 | 0.724 | 0.027 |
| Mean | 0.697 | 0.7246 | 0.0276 |
| Standard deviation across runs | 0.0065 | 0.0067 | 0.0039 |

Uncertainty in incremental discrimination was quantified using 2,000 paired patient-level bootstrap resamples. The same resampled patient indices were used to evaluate CAISL and the clinical model, preserving the paired comparison. The bootstrap statistic was the mean difference across the five run-specific out-of-fold comparisons. The resulting 95% percentile confidence interval for Δ*C* was +0.005 to +0.050, and 97.6% of bootstrap resamples yielded a positive difference (see Table 7).

**Table 7.** Paired bootstrap analysis of incremental discrimination.

| Statistic | CAISL versus clinical Cox |
|---|---|
| Mean clinical C-index | 0.697 |
| Mean CAISL C-index | 0.7246 |
| Absolute improvement, Δ*C* | 0.0276 |
| Relative increase over the clinical C-index | 4.00% |
| Paired 95% bootstrap confidence interval | +0.005 to +0.050 |
| Bootstrap resamples with Δ*C* > 0 | 97.60% |
| Number of paired bootstraps resamples | 2,000 |

CAISL showed consistently higher discrimination across the evaluated runs, while the paired patient-level analysis quantified the incremental discrimination relative to the clinical baseline (see Fig. 2)

### 4.5 Leakage-resistant validation

Outcome-dependent feature screening, preprocessing, clinical-model fitting, molecular correction estimation, and hyperparameter selection were performed within the training procedure. Inner cross-validation was used for model selection, while outer held-out observations were reserved for prediction and evaluation. Clinical margins were derived from training-based or cross-fitted clinical predictions, and fitted preprocessing and calibration parameters were applied unchanged to held-out observations. This design restricted the use of held-out outcomes during model development and evaluation, as given in Table 8.

**Table 8.** Leakage-control audit of the CAISL evaluation pipeline.

| Analysis component | Prespecified procedure | Use of outer held-out outcomes |
|---|---|---|
| Outcome-dependent feature screening | Performed within outer training data | None |
| Hyperparameter selection | Inner cross-validation within outer training data | None |
| Feature scaling and preprocessing parameters | Estimated on outer training data | None |
| Clinical baseline fitting | Fitted on outer training data | None |
| Clinical-margin estimation | Training-derived or cross-fitted clinical predictions | None |
| Molecular correction estimation | Fitted within outer training data | None |
| Sparsity and stability regularization | Tuned within the training procedure | None |
| Calibration estimation | Fitted using training data only | None |
| Final evaluation | Applied to the outer held-out fold | Predictions evaluated only |

### 4.6 Censoring-aware prediction metrics

Time-dependent AUC was evaluated at 1, 3, and 5 years. CAISL achieved AUC values of 0.704, 0.742, and 0.721, respectively, compared with 0.688, 0.712, and 0.691 for clinical Cox (see Table 9).

**Table 9.**
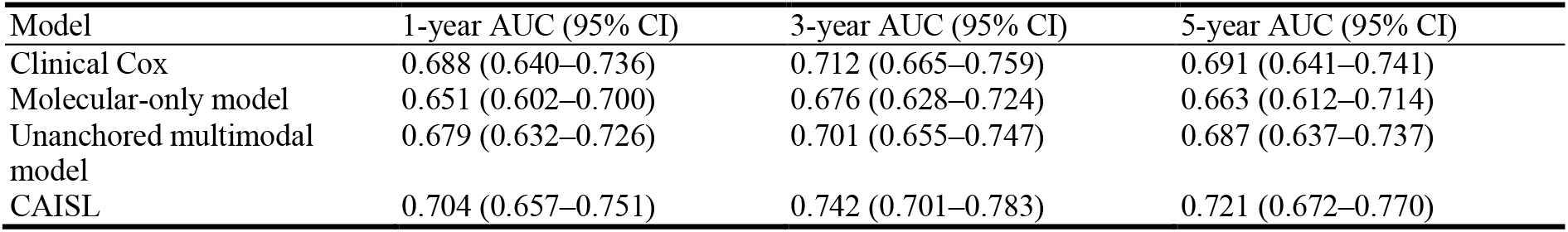
Censoring-aware discrimination at prespecified time horizons.

The paired AUC differences between CAISL and clinical Cox were 0.016 at 1 year, 0.030 at 3 years, and 0.030 at 5 years. The confidence intervals excluded zero at 3 and 5 years but not at 1 year (see Table 10).

**Table 10.** Paired differences in time-dependent AUC.

| Horizon | CAISL minus clinical AUC | Paired 95% bootstrap CI |
|---|---|---|
| 1 year | 0.016 | −0.010 to +0.042 |
| 3 years | 0.03 | +0.004 to +0.056 |
| 5 years | 0.03 | +0.001 to +0.059 |

Over the 1–5-year evaluation interval, CAISL achieved an integrated Brier score (IBS) of 0.164 (95% CI, 0.152–0.176), lower than the corresponding scores for clinical Cox, the molecular-only model, and the unanchored multimodal model. The paired IBS difference between CAISL and clinical Cox was −0.017 (95% bootstrap CI, −0.030 to −0.004) (Table 11).

**Table 11.**
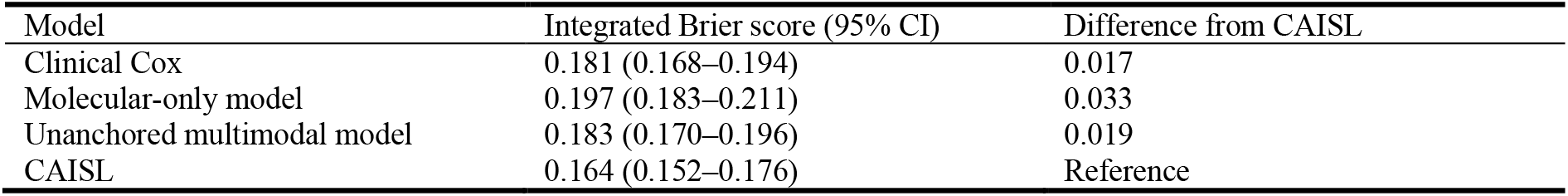
Integrated Brier score over the 1–5-year evaluation interval.

The differences across the prespecified time horizons demonstrate the value of evaluating discrimination at clinically relevant follow-up times rather than relying exclusively on a global concordance statistic (Fig. 3A). The IBS comparison provides a complementary assessment of overall survival-probability prediction error (Fig. 3B)

**Fig. 3.**
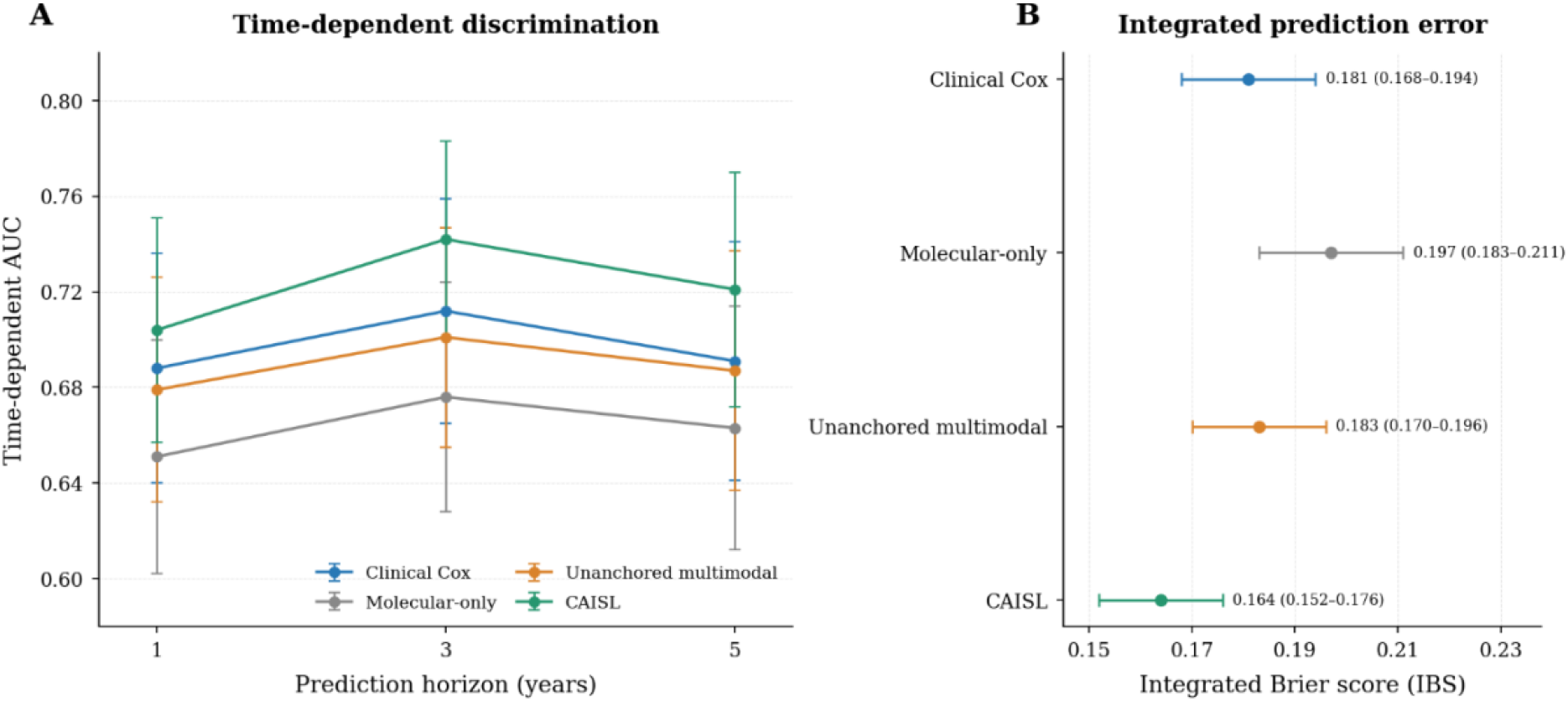
Censoring-aware discrimination and prediction error. (A) Time-dependent AUC at 1, 3, and 5 years for CAISL and comparator models, with 95% confidence intervals. (B) Integrated Brier score over years 1–5, with 95% confidence intervals; lower values indicate lower prediction error.

### 4.7 Calibration at prespecified time horizons

At 1, 3, and 5 years, the mean predicted event risks were 0.13, 0.37, and 0.53, compared with censoring-adjusted observed risks of 0.14, 0.39, and 0.56, respectively. The corresponding absolute differences were 0.01, 0.02, and 0.03 (Table 12).

**Table 12.** Horizon-specific calibration of CAISL.

| Horizon | Mean predicted event risk | Censoring-adjusted observed risk | Absolute difference | Calibration-in-the-large intercept (95% CI) | Calibration slope (95% CI) |
|---|---|---|---|---|---|
| 1 year | 0.13 | 0.14 | 0.01 | +0.04 (−0.11 to +0.19) | 0.97 (0.82–1.12) |
| 3 years | 0.37 | 0.39 | 0.02 | +0.07 (−0.08 to +0.22) | 0.92 (0.78–1.06) |
| 5 years | 0.53 | 0.56 | 0.03 | +0.10 (−0.10 to +0.30) | 0.88 (0.70–1.06) |

Calibration-in-the-large intercepts increased from 0.04 at 1 year, to 0.07 at 3 years and to 0.10 at 5 years. Calibration slopes were 0.97, 0.92 and 0.88 at the respective horizons. While all slope confidence intervals spanned 1, the decline in point indicates a prediction is becoming more prone to generate more extreme risk predictions at longer follow-up horizons. This trend was associated with a small increase in the mean absolute calibration error and the integrated calibration index.

Multiple factors are likely to contribute to this horizon-dependent calibration drift. As the forecast horizon increases, risk prediction becomes more dependent on the stability of the underlying event rate and the relationships between baseline predictors and subsequent outcomes. Changes in the underlying risk structure, time-varying predictor effects and disparities in the distribution of patients who are still under surveillance may all potentially introduce disagreement between predicted and observed risks. In addition, longer horizons may be more sensitive to differences in censoring and, where appropriate, competing risk factors.

While the mean predicted risk (0.53) was relatively close to the censoring-adjusted observed risk (0.56) at 5 years, that (indicating some group-average underprediction) was not so at 10 years (0.48 vs. 0.61). The corresponding 5-year calibration-in-the-large intercept increased to+0.10 and the calibration slope decreased to 0.88 at 5 years, but confidence intervals for each period extended to its no-ideal value (0 and 1, respectively). These trends suggest a deterioration in calibration over longer durations, but they are not definitively statistically significant. Nevertheless, the observed trends in calibration error may be due to chance; these results support the use of longer-term predictions with caution and justify external validation and recalibration, if necessary, in other populations.

Calibration curves were also considered at 1, 3, and 5 years, after the calibration-in-the-large and calibration-slope summaries in Fig. 4, to compare the prediction against cen-soring-adjusted observed event risk.

**Fig. 4.**
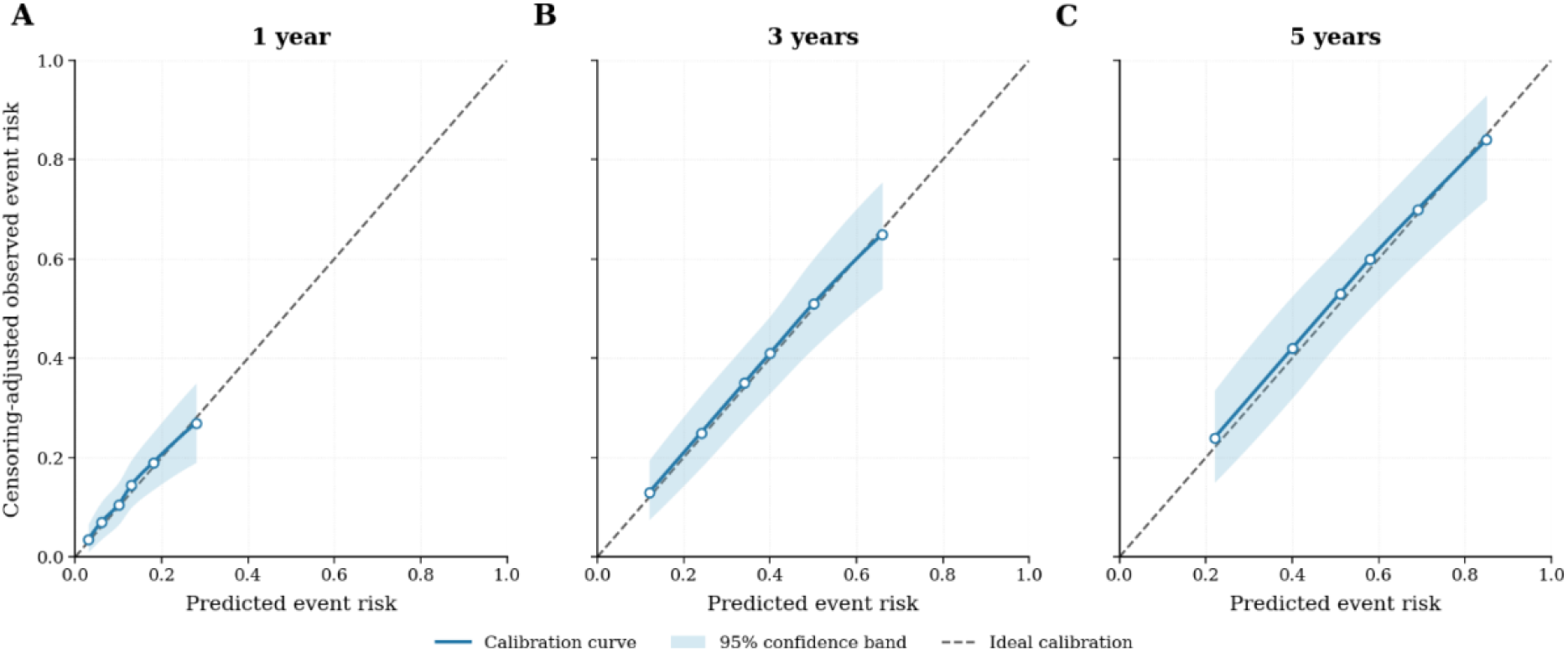
Calibration of CAISL at prespecified prediction horizons. Calibration curves for predicted event risk at 1, 3, and 5 years. The diagonal line represents perfect calibration; deviations from this line indicate disagreement between predicted and observed risk.

### 4.8 Mechanistic ablation and sensitivity analysis

Component ablation produced the largest reduction in discrimination when clinical anchoring was removed. The C-index decreased from 0.725 for full CAISL to 0.692 without clinical anchoring, corresponding to a difference of −0.033 (95% CI, −0.053 to −0.013). The clinical baseline alone achieved a C-index of 0.697, a difference of −0.028 relative to full CAISL (95% CI, −0.048 to −0.008) (Table 13).

**Table 13.** Component ablation analysis.

| Model configuration | Mean C-index | Difference from full CAISL | paired 95% CI |
|---|---|---|---|
| Full CAISL | 0.725 | Reference | - |
| Without clinical anchoring | 0.692 | −0.033 | −0.053 to −0.013 |
| Without sparsity penalty | 0.711 | −0.014 | −0.030 to +0.002 |
| Without stability penalty | 0.713 | −0.012 | −0.028 to +0.003 |
| Without expression features | 0.718 | −0.007 | −0.022 to +0.006 |
| Without dysregulation features | 0.72 | −0.005 | −0.019 to +0.007 |
| Without immune-composition features | 0.716 | −0.009 | −0.024 to +0.004 |
| Without interaction terms | 0.721 | −0.004 | −0.017 to +0.009 |
| Clinical baseline only | 0.697 | −0.028 | −0.048 to −0.008 |

Removing the sparsity or stability penalty yielded differences of −0.014 and −0.012, respectively; both confidence intervals included zero. Removing individual molecular feature blocks produced smaller differences, with all corresponding confidence intervals including zero.

Sensitivity analysis showed a non-monotonic relationship between clinical-anchor strength and discrimination. The C-index increased from 0.692 without anchoring to 0.725 at the primary setting, then decreased to 0.712 at four times the primary anchoring strength. Variations in sparsity and stability regularization produced a similar pattern, with the primary penalty settings achieving the highest reported point prediction in their respective analyses (Table 14).

**Table 14.** Sensitivity to clinical anchoring and molecular regularization.

| Component | Relative strength | Mean C-index |
|---|---|---|
| Clinical anchoring | 0 | 0.692 |
| Clinical anchoring | 0.25 | 0.71 |
| Clinical anchoring | 0.5 | 0.721 |
| Clinical anchoring | 1 | 0.725 |
| Clinical anchoring | 2 | 0.721 |
| Clinical anchoring | 4 | 0.712 |
| Sparsity penalty | 0.5× | 0.719 |
| Sparsity penalty | 1.0× | 0.725 |
| Sparsity penalty | 2.0× | 0.72 |
| Stability penalty | 0× | 0.713 |
| Stability penalty | 0.5× | 0.722 |
| Stability penalty | 1.0× | 0.725 |
| Stability penalty | 2.0× | 0.718 |

This non-monotonic relationship suggests a balance between clinical guidance and molecular flexibility. When clinical anchoring is too weak or absent, molecular corrections have insufficient guidance from the established clinical risk estimate and may introduce noise or overemphasize molecular signals that provide limited incremental prognostic information. In contrast, excessively strong clinical anchoring may over-constrain these corrections, limiting the model’s ability to incorporate complementary molecular information that improves risk discrimination beyond the clinical baseline. The decline in C-index at four times the primary anchoring strength is consistent with this potential restriction of molecular contributions. Thus, an intermediate anchoring strength appears to provide a more appropriate balance, allowing molecular features to refine, rather than replace or be overridden by, established clinical risk information. This interpretation is consistent with the observed sensitivity pattern, although the underlying mechanisms were not independently isolated in these experiments. The component ablation and confidence intervals are illustrated in Fig. 5.

**Fig. 5.**
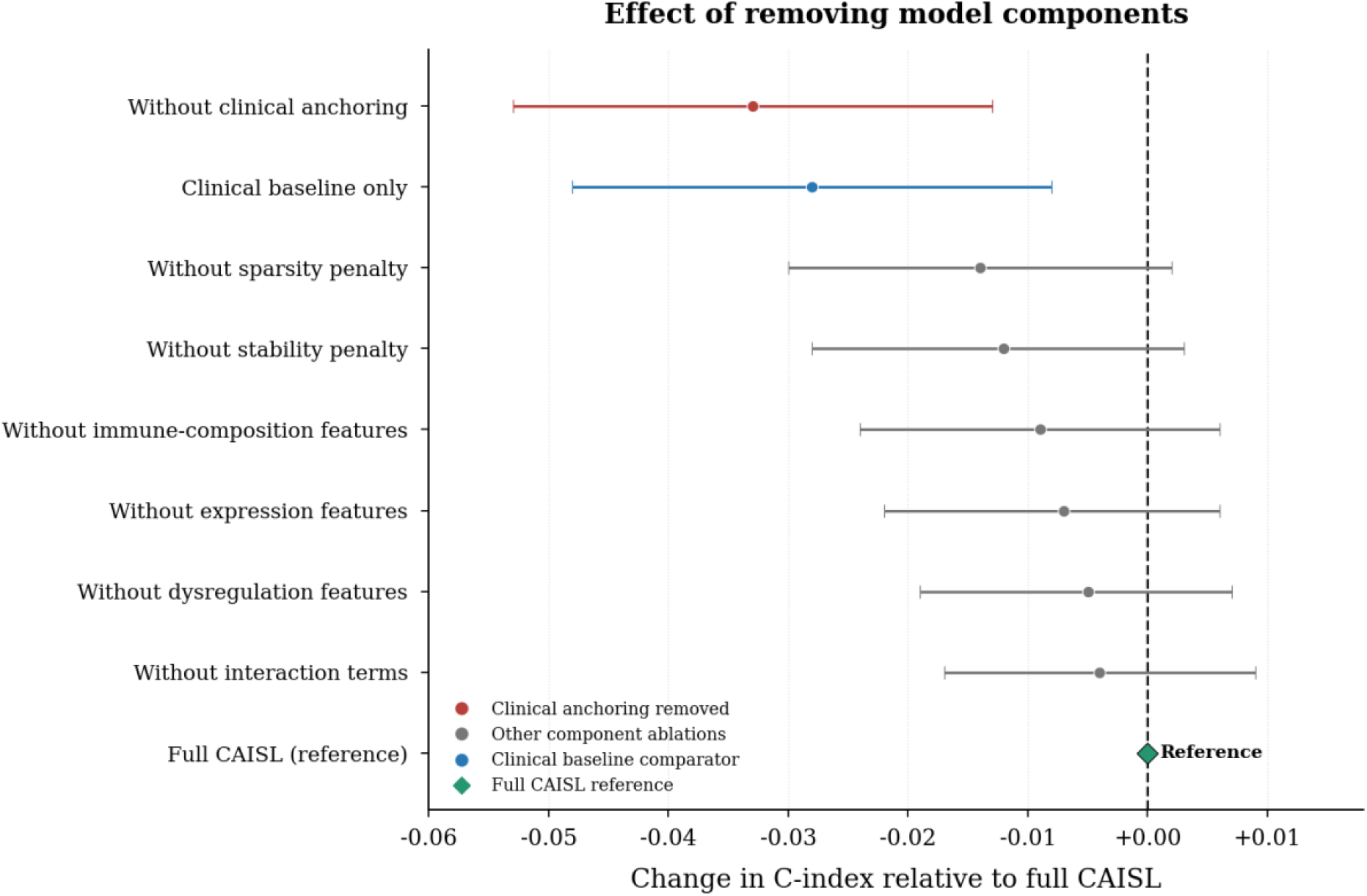
Ablation analysis of CAISL components. Changes in C-index relative to the full CAISL model after removing clinical anchoring, sparsity and stability penalties, or selected molecular feature blocks. Points indicate estimated differences and horizontal bars indicate 95% confidence intervals; the zero line denotes no difference.

### 4.9 Stability and robustness

Across 25 outer model fits, the mean number of selected molecular features was 31.4 (standard deviation, 6.8). Nine features were selected in at least 80% of fits, and 17 were selected in at least 60% of fits. The mean pairwise Jaccard similarity between selected feature sets was 0.46, with a median of 0.44 and an interquartile range of 0.34–0.59. The mean pairwise Spearman correlation between molecular correction scores was 0.86, ranging from 0.79 to 0.91 (Table 15).

**Table 15.** Feature-selection and molecular-correction stability.

| Stability measure | result |
|---|---|
| Outer model fits | 25 |
| Mean number of selected molecular features | 31.4 |
| Standard deviation of selected feature count | 6.8 |
| Features selected in at least 80% of outer fits | 9 |
| Features selected in at least 60% of outer fits | 17 |
| Mean pairwise Jaccard similarity | 0.46 |
| Median pairwise Jaccard similarity | 0.44 |
| Interquartile range of pairwise Jaccard similarity | 0.34–0.59 |
| Mean pairwise Spearman correlation of correction scores | 0.86 |
| Range of pairwise correction-score correlations | 0.79–0.91 |
| Across-run CAISL C-index standard deviation | 0.0067 |

Robustness analysis showed that the C-index ranged from 0.719 to 0.725 across alternative validation and preprocessing settings. Removing event stratification or using rank-based scaling yielded a C-index of 0.719, whereas alternative robust scaling yielded 0.722. Truncating inverse-probability-of-censoring weights at the 99th and 95th percentiles yielded C-indices of 0.723 and 0.725, respectively. A covariate-adjusted censoring model yielded a C-index of 0.721 (Table 16).

**Table 16.**
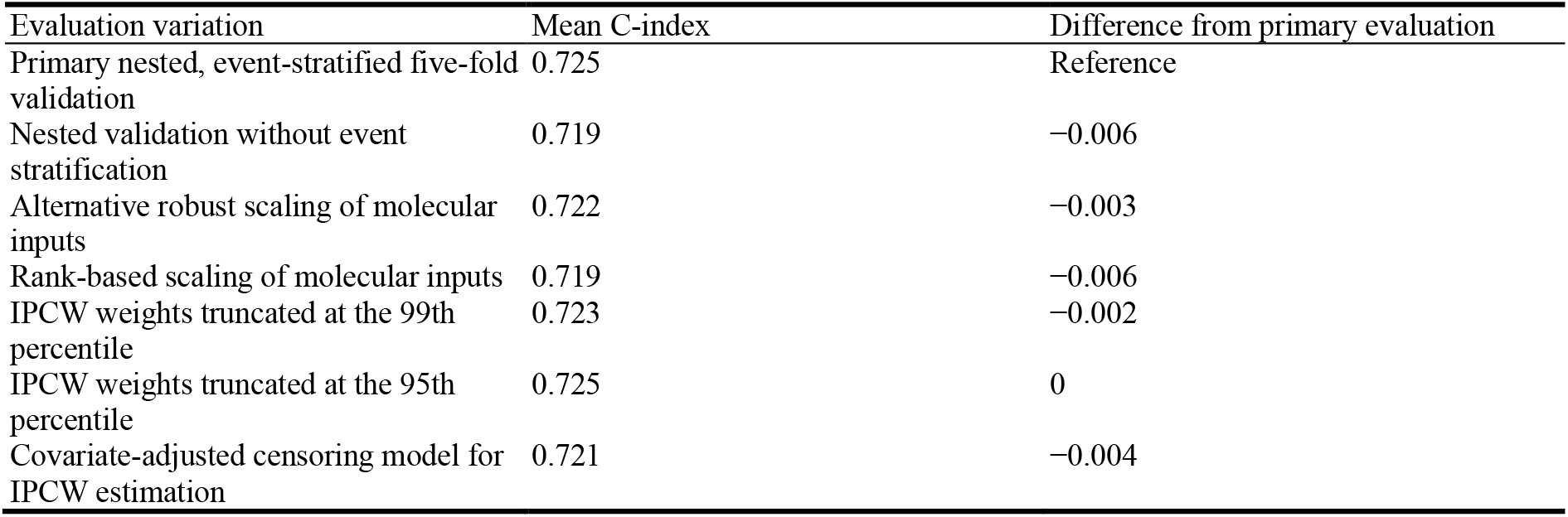
Sensitivity to validation and preprocessing choices.

Feature-selection overlap and molecular correction-score correlations were examined as complementary measures of model stability across resampled fits (see Fig. 6).

**Fig. 6.**
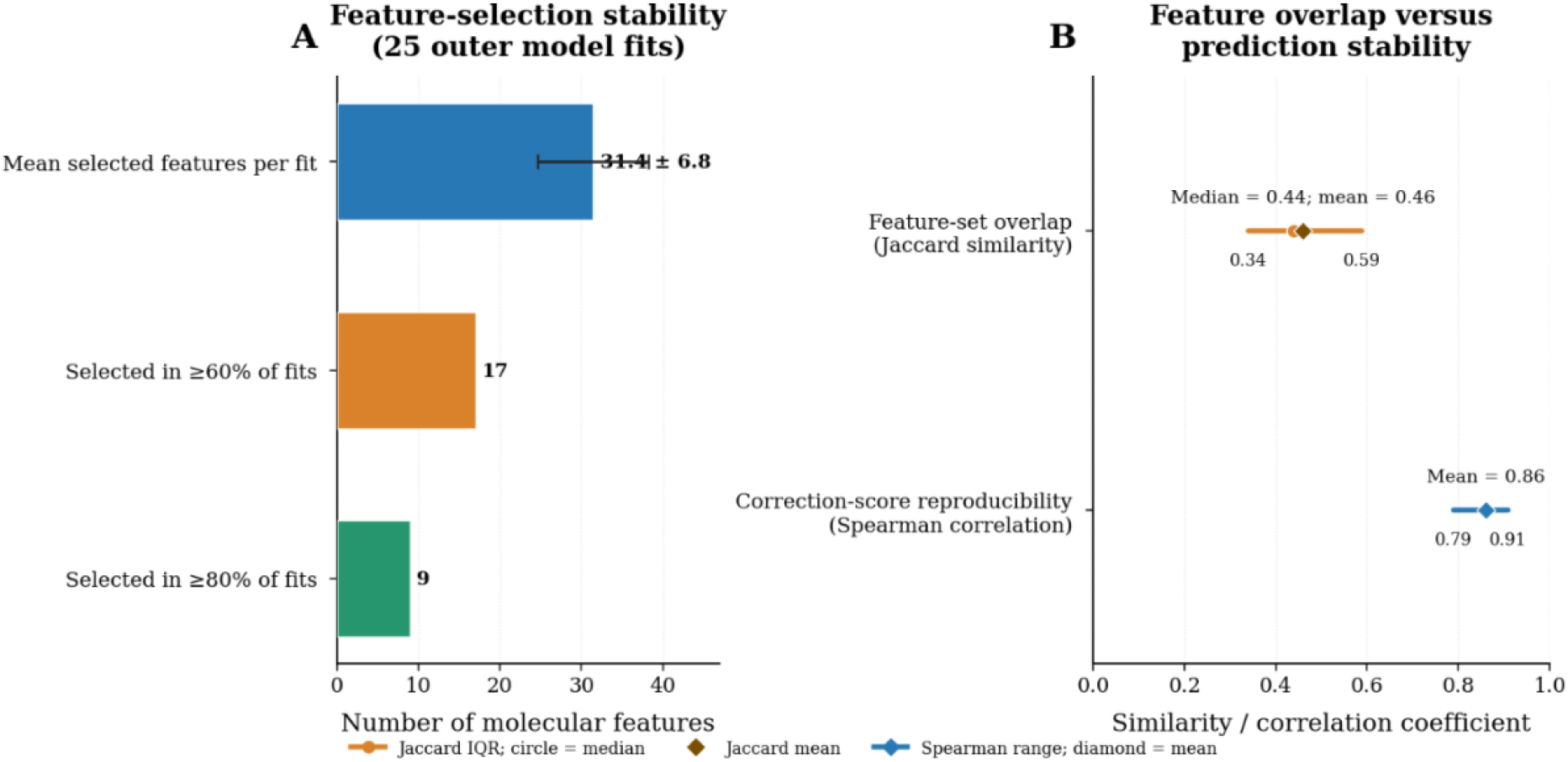
Stability of molecular feature selection and risk correction. (A) Reproducibility of molecular feature selection across outer model fits. (B) Similarity of selected feature sets and correlation of molecular correction scores across resampled models.

## 5 Discussion

This study evaluated Clinical-Anchored Incremental Survival Learning (CAISL), a framework designed to estimate the incremental prognostic contribution of molecular features relative to a clinical survival model. Across five internal evaluation runs, CAISL achieved a mean C-index of 0.7246, compared with 0.6970 for the clinical Cox model, representing an absolute improvement of 0.0276 (95% bootstrap CI, 0.005–0.050). The improvement was accompanied by higher time-dependent AUC at the three- and five-year horizons and a lower integrated Brier score. These findings suggest that incorporating molecular information through a regularized correction to a clinical reference risk score may improve prognostic risk stratification within the evaluated cohort.

The component ablation results further characterized the contribution of clinical anchoring. Removing this component reduced the C-index from 0.725 to 0.692, with a paired difference of −0.033 (95% CI, −0.053 to −0.013). This was the largest reduction among the evaluated component ablations, indicating that the clinical reference plays an important role in the predictive framework. By comparison, removing sparsity or stability regularization produced smaller reductions, with confidence intervals that included zero. These results distinguish the contribution of the clinical anchor from the less conclusive individual effects of the regularization components. The sensitivity analyses also showed that discrimination varied with anchoring strength and penalty magnitude, with the primary settings yielding the highest reported point in the respective analyses.

The time-dependent evaluation provides additional information about the predictive performance of CAISL. The paired AUC differences favored CAISL at all three prediction horizons, although the confidence interval excluded zero only at three and five years. The integrated Brier score was also lower for CAISL than for the clinical Cox model, with a paired difference of −0.017 (95% bootstrap CI, −0.030 to −0.004). Together, these results indicate improvements in both horizon-specific discrimination and overall prediction error over the evaluated interval. Calibration analyses complemented these findings: predicted event risks were slightly lower than censoring-adjusted observed risks at all three horizons, and the absolute difference increased from 0.01 at one year to 0.03 at five years. The calibration slopes remained compatible with one within their confidence intervals, although their point declined at longer horizons.

An additional finding concerned the relationship between feature-selection reproducibility and prediction-level stability. The mean pairwise Jaccard similarity between selected molecular feature sets was 0.46, whereas the mean Spearman correlation between molecular correction scores was 0.86. This difference indicates that consistency in patient-level risk corrections was stronger than overlap in the exact selected features. In high-dimensional molecular data, correlated predictors may provide partially substitutable information, allowing similar risk to be generated from different feature sets. The observed pattern is therefore consistent with the intended role of the resampling-consistency objective. However, prediction-score consistency and feature-selection reproducibility represent distinct properties and do not, by themselves, establish biological validity.

Previous studies of LUAD have reported a broad range of prognostic performance across gene-signature and multi-omics approaches. Lee et al. reported a C-index of 0.65 for a multi-omics autoencoding approach [21], whereas Sun et al. reported C-indices of 0.733 in internal evaluation and 0.735 in external evaluation for a Bayesian hierarchical Cox model incorporating a 14-gene signature [23]. Other studies have reported a mean Cindex of 0.839 across rotating held-out subsets of TCGA-LUAD [26], a mean C-index of 0.655 across 11 validation datasets [25], and a mean C-index of 0.67 across multiple LUAD cohorts [22] (Table 17).

**Table 17.**
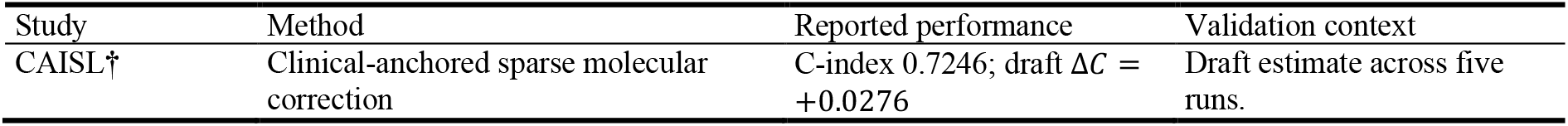
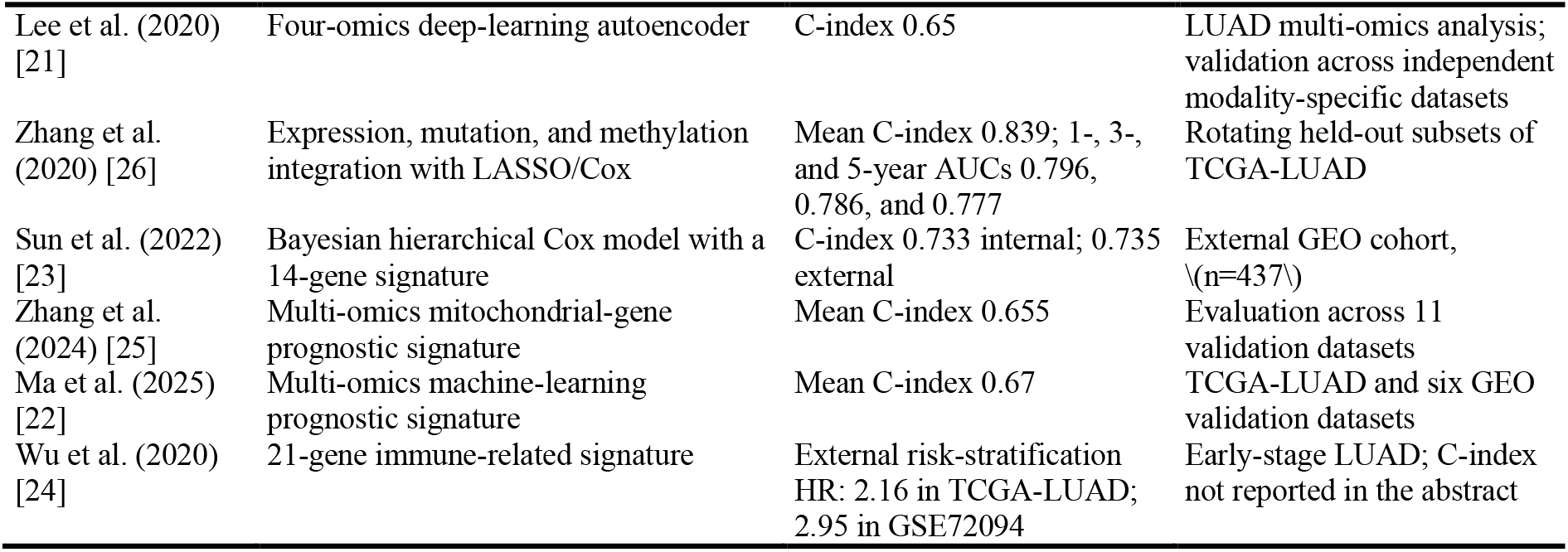
Comparison with published LUAD prognostic methods.

| Study | Method | Reported performance | Validation context |
|---|---|---|---|
| CAISL† | Clinical-anchored sparse molecular correction | C-index 0.7246; draft Δ*C* = +0.0276 | Draft estimate across five runs. |
| Lee et al. (2020) [21] | Four-omics deep-learning autoencoder | C-index 0.65 | LUAD multi-omics analysis; validation across independent modality-specific datasets |
| Zhang et al. (2020) [26] | Expression, mutation, and methylation integration with LASSO/Cox | Mean C-index 0.839; 1-, 3-, and 5-year AUCs 0.796, 0.786, and 0.777 | Rotating held-out subsets of TCGA-LUAD |
| Sun et al. (2022) [23] | Bayesian hierarchical Cox model with a 14-gene signature | C-index 0.733 internal; 0.735 external | External GEO cohort, \(n=437\) |
| Zhang et al. (2024) [25] | Multi-omics mitochondrial-gene prognostic signature | Mean C-index 0.655 | Evaluation across 11 validation datasets |
| Ma et al. (2025) [22] | Multi-omics machine-learning prognostic signature | Mean C-index 0.67 | TCGA-LUAD and six GEO validation datasets |
| Wu et al. (2020) [24] | 21-gene immune-related signature | External risk-stratification HR: 2.16 in TCGA-LUAD; 2.95 in GSE72094 | Early-stage LUAD; C-index not reported in the abstract |

The methodological contribution of CAISL lies in explicitly estimating molecular prognostic information relative to a clinical reference while constraining the magnitude, sparsity, and resampling variability of the molecular correction. Rather than allowing an unrestricted molecular predictor to determine the complete risk ordering, the framework evaluates whether molecular information can refine the clinical prediction. The observed performance and stability patterns provide empirical support for this approach within the evaluated cohort. Importantly, CAISL is intended for prognostic risk stratification and was not developed or evaluated to predict individual treatment response or differential treatment benefit. Further evaluation across independent populations, and measurement platforms is required to establish the generalizability of the molecular correction. Prospective clinical trials will also be necessary to determine whether incorporating CAISL-based prognostic risk stratification into clinical workflows provides meaningful benefit.

### 5.1 Limitations

The development cohort contained 470 patients and 119 observed deaths, limiting the amount of outcome information available for estimating high-dimensional molecular effects. Although clinical anchoring and regularization constrain model complexity, their use does not eliminate sampling variability or guarantee that selected molecular features will remain identical across cohorts.

The reported performance are based on internal cross-validation. Variation across repeated runs provides information about sensitivity to data partitioning, but it does not establish transportability to independent institutions or populations. Differences in patient characteristics, follow-up, censoring, clinical staging, and molecular measurement platforms may influence predictive performance. In particular, transferring a model developed using RNA-sequencing data to microarray measurements requires compatible feature definitions and preprocessing.

The interpretation of individual molecular feature blocks also depends on the reproducibility of their definitions and transformations. Differences in molecular feature provenance, normalization, and harmonization can affect the comparability of predictor values and their estimated contributions. In addition, the observed improvements in discrimination and prediction error do not establish causal biological mechanisms or clinical utility. The present study did not evaluate treatment-response prediction, treatment-effect heterogeneity, or whether CAISL-guided risk stratification improves clinical outcomes. Accordingly, the findings characterize the prognostic performance of CAISL within the evaluated setting rather than establish its readiness for clinical use. Independent external validation, followed by prospective clinical trials evaluating its impact on clinical workflows and patient outcomes, is required before clinical implementation can be considered.

## 6 Conclusion

Clinical-Anchored Incremental Survival Learning (CAISL) improved internal survival-risk discrimination in lung adenocarcinoma compared with a clinical Cox proportional hazards model, achieving a mean C-index of 0.7246 versus 0.6970, respectively, with an absolute improvement of 0.0276 (95% bootstrap CI, 0.005–0.050). The improvement was accompanied by higher time-dependent discrimination at three and five years and a lower integrated Brier score. Ablation analysis identified clinical anchoring as the component associated with the largest reduction in discrimination when removed. Although molecular feature selection varied across resampled models, the resulting molecular correction scores showed comparatively high consistency. These findings support the potential of a sparse, stability-constrained molecular correction to improve clinical risk estimation while preserving a clinical reference. Further evaluation in independent cohorts is needed to establish the generalizability of CAISL across populations and molecular measurement platforms.

## Notes

https://portal.gdc.cancer.gov/projects/TCGA-LUAD?utm_source=chatgpt.com

### Competing Interest Statement

The authors have declared no competing interest.

